# EZpred: improving deep learning-based enzyme function prediction using unlabeled sequence homologs

**DOI:** 10.1101/2025.07.09.663945

**Authors:** Chengxin Zhang, Quancheng Liu, Lydia Freddolino

## Abstract

Features extracted from sequence homologs significantly enhance the accuracy of deep learning-based protein structure prediction. Indeed, models such as AlphaFold, which extracts features from sequence homologs, generally produce more accurate protein structures compared to single sequence-based methods like ESMfold. In contrast, features from sequence homologs are seldom employed for deep learning-based protein function prediction. Although a small number of models also incorporate function labels from sequence homologs, they cannot utilize features extracted from sequence homologs that lack function labels. To address this gap, we propose EZpred, which is the first deep learning model to use unlabeled sequence homologs for protein function prediction. Starting with the target sequence and homologs identified by MMseqs2, EZpred extracts sequence features using the ESMC protein language model. These features are then fed into a deep learning model to predict the Enzyme Commission (EC) numbers of the target protein. For 753 enzymes, the F1-score of EZpred EC number prediction is 4% higher than a similar model that does not use sequence homologs and at least 10% higher that state-of-the-art EC number prediction models. These results demonstrate the strong positive impact of sequence homologs in deep learning-based enzyme function prediction.

**Significance Statement:** Multiple sequence alignment (MSA) of homologous sequences is the most important source of input feature for deep learning-based protein structure prediction. Yet, this kind of feature is rarely used for protein function prediction. We propose the first deep learning model that significantly improves protein function prediction using features from sequence homologs without functional labels. This shows the utility of an important set of features that are often overlooked by previous studies.

## Introduction

In the past five years, the development of deep learning models for high accuracy protein structure prediction is the most important breakthrough in computational biology. Notable successful structure prediction models include AlphaFold (1, 2), RoseTTAFold (3), and D-I-TASSER (4). In all these models, the target protein sequence is searched through one or more sequence databases to identify sequence homologs. Although these homologs do not have known structures, their alignment is still used to extract input features for the deep learning models. As the entry point of protein structure prediction, homologous sequence search and alignment have a strong impact on the accuracy of predicted protein structure models (5, 6). On the other hand, single sequence-based protein structure models such as ESMfold (7) that extracts features from the target sequence alone generally have worse accuracy than models that use homologous sequence alignments, as shown by recent Critical Assessment of Structure Prediction 15 (CASP15) challenge (8).

Although the usage of homologous sequences has been highly successful in deep learning-based protein structure prediction, similar success has not been replicated in the prediction of protein functions, especially Gene Ontology (GO) terms and Enzyme Commission (EC) numbers. Traditionally, protein function prediction was achieved by transferring function annotations from homologous detected by sequence and structure searches. For example, GOtcha (9), Blast2GO (10), ConFunc (11), PFP (12), and CombFunc (13), and GoFDR (14) extracts GO terms from templates detected by sequence search performed using BLASTp (15), PSI-BLAST (15) or DIAMOND (16). Our recent study shows that when using optimized sequence search parameters, BLASTp, DIAMOND and MMseqs2 (17) can all achieve comparable performance (18). Apart from sequence templates, COFACTOR (19, 20), StarFunc (21) and ProFunc use function annotations from templates detected by structure alignment performed using TM-align (22), Foldseek (23), SSM (24) and Jess (25). Apart from sequence templates and structure templates, other template information such as protein-protein interaction partners and protein domain families is also used for function prediction (26-30).

While traditional protein function annotation tools are mainly based on template search, most deep learning-based methods for protein function prediction extracts features solely from the target protein sequence. To represent the target protein sequence, some models such as DeepGO (31), DeepGOplus (32) and ProteInfer (33) use one-hot encoding, while other models such as ATGO+ (34), AnnoPRO (35), DeepGO-SE (36) and CLEAN (37) uses protein language models such as ESM (7, 38) or ProtT5(39). A small number of deep learning models such as SPROF-GO (40) and GraphEC (41) also use function annotations from sequence homologs to improve prediction accuracy.

When using sequence homologs for function prediction, both traditional template-based tools and deep learning methods only use sequence templates with known functions. This is different from deep learning-based protein structure prediction, where sequence homologs without known structures play an important role.

To enclose this gap, we propose EZpred, the first deep learning model that improves function predictions using sequence homologs without known function labels. It is specifically designed to predict enzyme functions in the form of EC numbers. EZpred searches the target enzyme sequence through a sequence database by MMseqs2 to detect enzyme homologs. Sequence features of the target protein and enzyme homologs are derived from the last three layers of the ESMC (42) protein language model. These features are average-pooled along the sequence and concatenated together. The concatenated features are fed into a multi-layer perception (MLP) network to predict EC numbers. Meanwhile, the target protein sequence and structure are used to search template database using MMseqs2 and Foldseek. EC numbers derived from templates are then combined with deep learning-based prediction to derive the final prediction (**Figure 1**). EZpred is available both as a webserver (https://seq2fun.dcmb.med.umich.edu/EZpred/) and as a source code distribution downloadable from that website.

**Figure 1.**
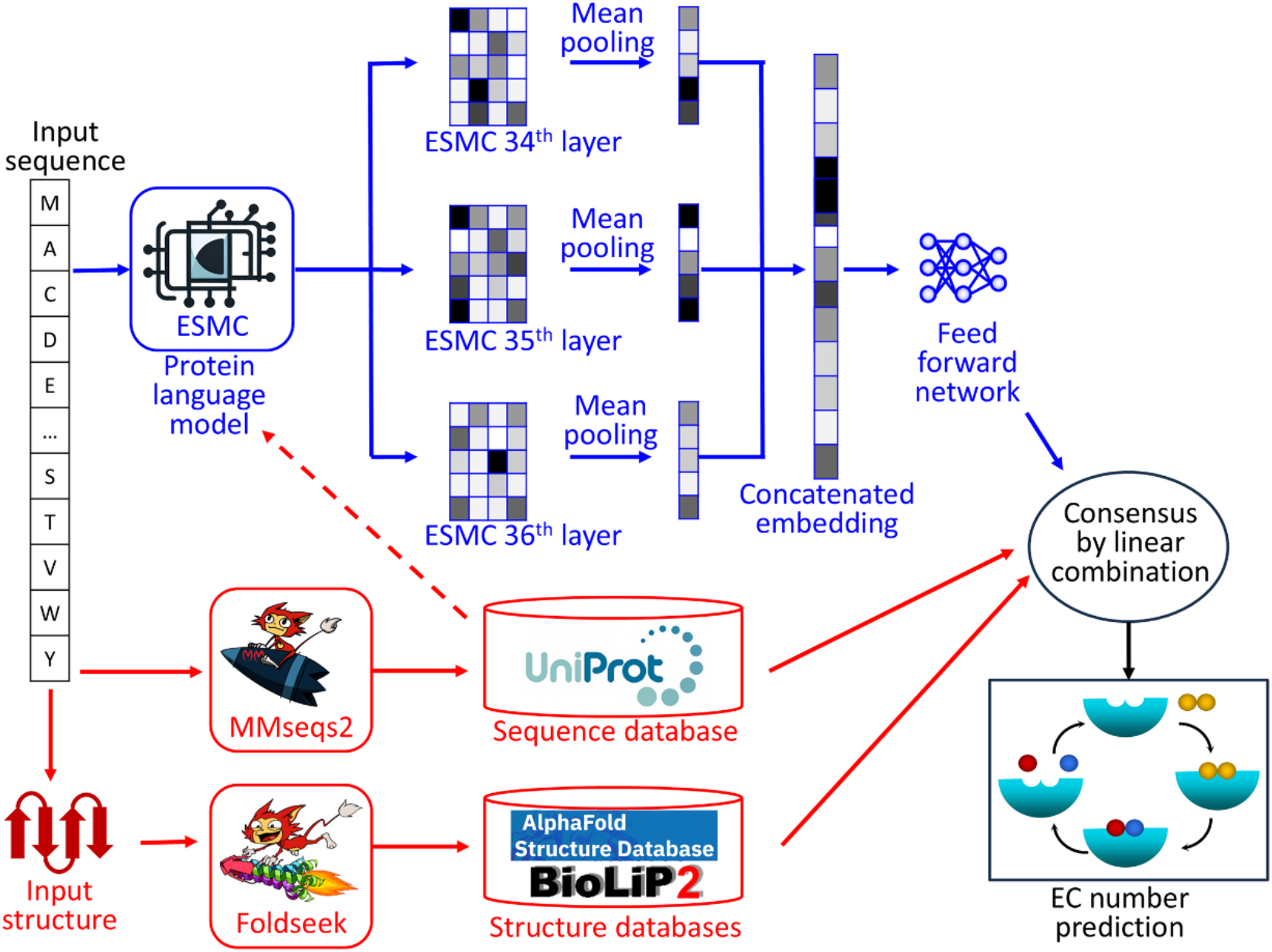
The EZpred pipeline. Blue indicates the deep learning (DL) part of EZpred, while red indicates the template-based part of EZpred.

## Results

### Dataset

EZpred was trained on 30592 enzymes curated from UniProt release 2023_02. To collect enzymes, we extract the EC number annotations based on the “DE”, “CC” and “DR BRENDA” fields of the plain text format UniProt data. Additionally, if the UniProt protein does not have EC number annotation but has Rhea (43) reaction annotation, we convert the Rhea ID to EC number using the rhea2ec mapping (https://ftp.expasy.org/databases/rhea/tsv/rhea2ec.tsv). Since these EC number annotations may come from either experiments or computational studies, we extract the subset of experimental EC number annotations based on the Evidence and Conclusion Ontology (ECO) term associated with the EC number or Rhea ID annotation, where ECO:0000305 and ECO:0000269 indicate experimental annotations. Additionally, if the UniProt protein has GO annotations that can be converted to EC number using the ec2go mapping (http://release.geneontology.org/2024-04-24/ontology/external2go/ec2go), and the GO term is annotated with an experimental evidence code (EXP, IDA, IPI, IMP, IGI, IEP, HTP, HDA, HMP, HGI, HEP, TAS, or IC), we will also collect this experimental annotation. These proteins are clustered by CD-HIT (44) using 60% sequence identity cutoff. We then partition sequence clusters into 5 folds, stratified by EC labels. Hyperparameters of EZpred are optimized by 5-fold cross validation.

Based on the same UniProt release, we also collect non-enzymes as the negative training set. To generate this dataset, we collect all UniProt proteins with at least one Molecular Function (MF) GO term annotated with an experimental evidence code. The following GO terms are excluded as they are too general: GO:0046872 “metal ion binding” as well as its child terms, GO:0005524 “ATP binding”, GO:0042802 “identical protein binding”, GO:0005515 “protein binding”, GO:0051260 “protein homooligomerization”, GO:0003676 “nucleic acid binding”. For the remaining proteins with at least one specific MF GO term annotation, we exclude enzymes that are annotated with EC numbers or any child terms of GO terms GO:0003824 “catalytic activity” or GO:0022857 “transmembrane transporter activity”. This results in 23354 non-enzyme proteins. This negative training set is also partitioned into 5-folds using a similar approach to the enzyme training set.

To generate an independent test set, we collect 753 enzymes that lack EC number annotation in UniProt release 2023_02 but have new EC number annotation in UniProt release 2024_02. We also include 625 negative test proteins that have new experimental MF GO term annotations to confirm non-enzyme activity in UniProt release 2023_02 but without such GO terms in UniProt release 2023_02.

### Overall Performance

The performance of EZpred and the pretrained models of 8 existing machine learning-based EC number prediction methods on 753 enzymes are evaluated by the F1-scores of predicting the first digit, the first two digits, the first three digits and all four digits of the EC number annotation. Here, F1-score is the harmonic average of precision and recall; a higher F1-score indicates more accurate prediction. Among the existing EC number predictors, ProteInfer (33), ECPICK (45) and ECPred (46), DeepEC (47) extract features from the raw protein sequence; CLEAN (37), and DeepECtransformer (48) use protein language models (39, 49) to extract features from the protein sequence. While these 7 methods only use the protein sequence for feature extraction, GraphEC (41) and CLEAN-Contact (50) use both the 3D structure (represented by a contact map) and the protein sequence (featurized by protein language model) to extract the features.

Overall, apart from DeepEC, all benchmarked methods can relatively accurately predict the first three digits of EC numbers with F1-score ≥0.646, where EZpred achieves the highest accuracy (F1-score=0.889, **Figure 2**). On the other hand, predicting all four digits EC number is much more challenging, where the F1-score ranging from 0.788 (EZpred) to 0.432 (DeepEC). The F1-score of EZpred is 27% and 32% higher than CLEAN and GraphEC, which are the two most accurate methods. While the template- and deep learning-based components of EZpred achieve a similar performance (F1-score=0.785), combining these two components results in an improvement of 0.4%. The main improvement is for the subset of hard targets. This improvement is modest, probably because both components have already achieved high performance, making further improvement difficult.

**Figure 2.**
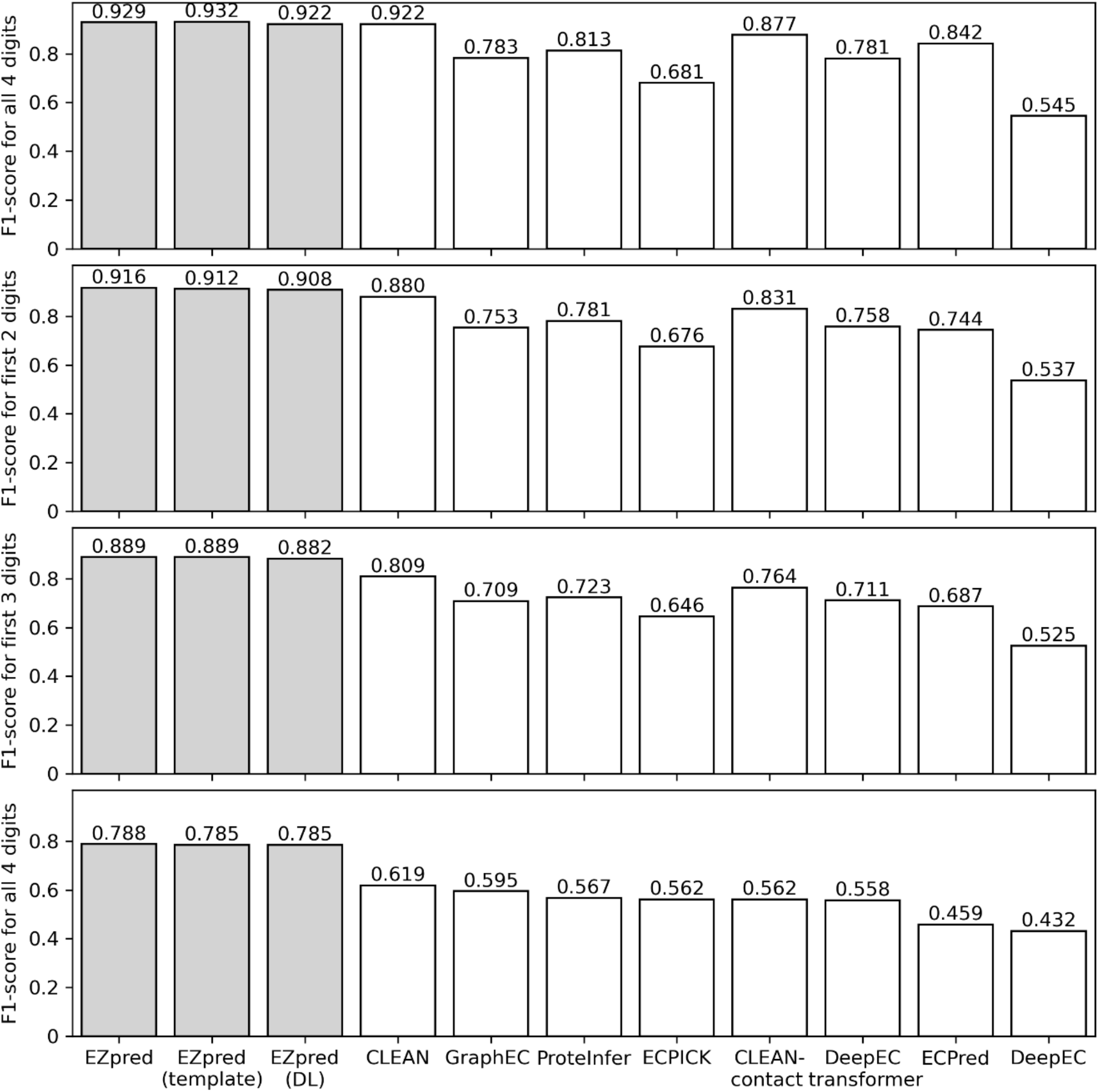
F1-score of EC number prediction for 753 test enzymes. EZpred (template) and EZpred (DL) indicate the template- and deep learning-based components of EZpred, respectively. Different methods are sorted in descending order of F1-score for all 4 EC number digits. Shaded bars indicate EZpred or its component pipelines.

To further investigate the impact of sequence similarity on model performance, we partition the test set into “easy”, “medium”, and “hard” target sets, where their maximum sequence identity to the most similar training enzyme is in the range of 0.6∼1, 0.3∼0.6 and 0∼0.3, respectively (**Figure 3**). The performance of all tested methods is worse for targets with lower sequence similarity. For all three sets of targets, EZpred as well as its component methods outperform existing methods. When predicting all 4 digits of EC number, although the deep learning-based component of EZpred slightly outperforms the template-based component for easy and medium targets, the former is worse than the latter for hard targets (F1-score=0.598 and 0.638, respectively). This is probably because the deep learning-based component only uses sequence information while the template-based component also uses the tertiary structure, which allows more sensitive function detection by the latter component for challenging targets.

**Figure 3.**
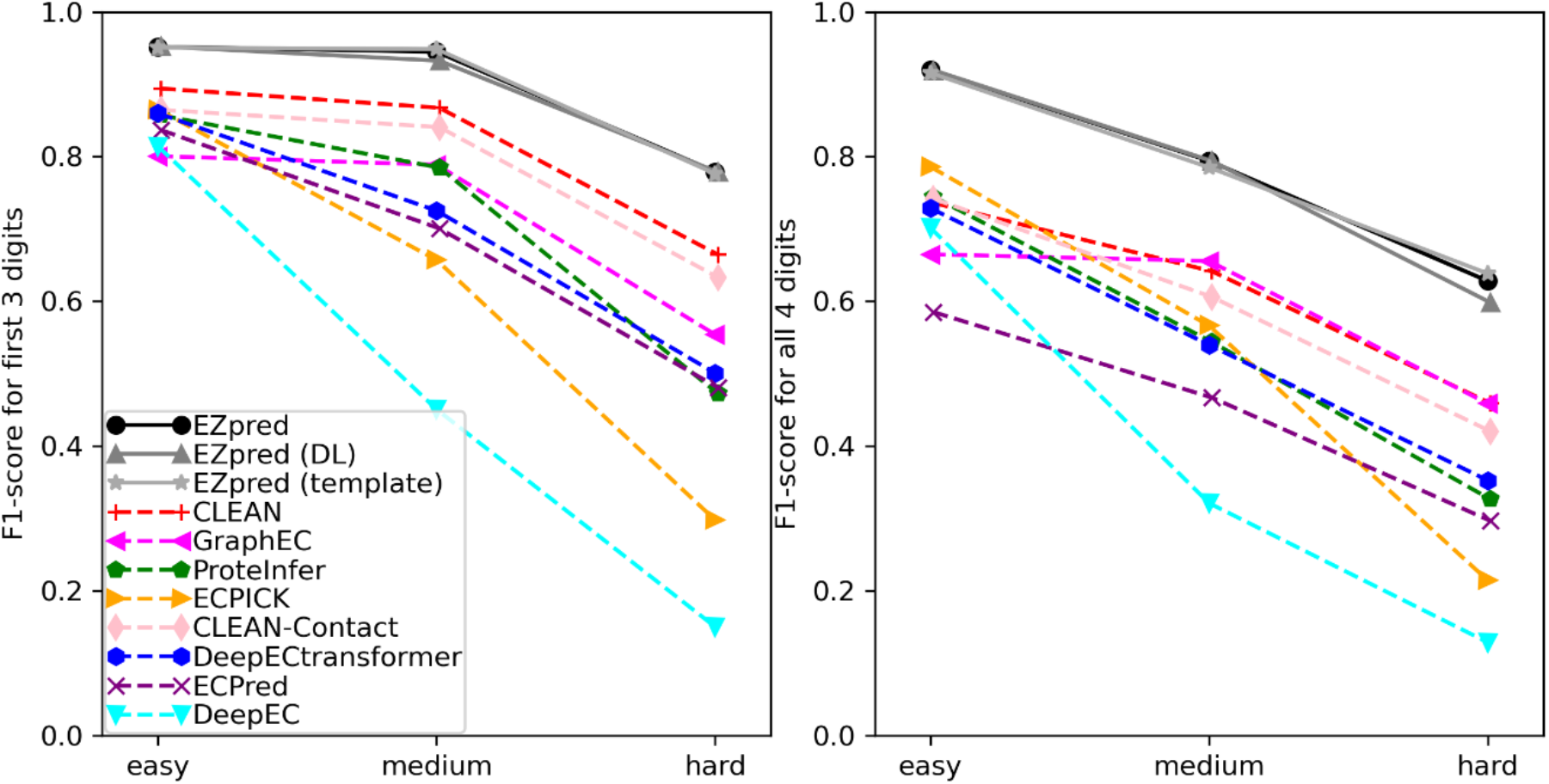
Performance of EZpred and existing methods for predicting the first 3 and all 4 digits of EC numbers on 252 “easy” targets, 248 “medium” targets and 253 “hard” targets among the 753 testing enzymes.

### Enzyme vs non-enzyme prediction

In many real case scenarios, we do not know whether the input protein is an enzyme or non-enzyme. Therefore, a practical enzyme function predictor should be able to differentiate enzymes versus non-enzymes. The deep learning-based component of EZpred has a separate label (EC number 0.-.-.-) for non-enzymes. DeepECtransformer and ECPred also predict similar non-enzyme label (“None” for DeepECtransformer and “non Enzyme” for ECPred). We compare the performance of EZpred, DeepECtransformer and ECPred in differentiating enzymes from non-enzymes. On the full testing dataset containing 753 enzymes and 625 non-enzymes, the F1-score for predicting whether a target protein is an enzyme is 0.911, 0.568 and 0.780 for EZpred, DeepECtransformer and ECPred, respectively. This shows EZpred is much more sensitive in identifying enzymes than existing methods.

### Ablation study

EZpred is different from many other deep learning-based protein function prediction models in the choices of different hyperparameters and feature representations. To quantify the contributions of these choices, we perform a set of controlled experiments. First, many methods exclude rare function labels during training. For example, InterLabelGO (51) excludes GO terms with less than 10 training proteins; DeepECtransformer excludes EC numbers with less than 100 training enzymes. On the other hand, EZpred includes any EC number in the training set, even if there is only one training protein associated with the EC number. Including all EC numbers results in 5% and 16% higher F1-score than the same model architecture including EC numbers with at least 5 or 10 training enzymes (**Table 1**). A probable reason that including rare function labels improves EZpred but not previous methods is our usage of a ranking-based loss function (See Methods). This ranking-based loss makes the learning of sparse labels effective, which may not be true for previous methods that use binary cross entropy-based loss.

**Table 1.** F1-score of predicting all 4 digits of EC number prediction by different versions of the deep learning component of EZpred on 753 test enzymes.

| Minimal number of training enzymes for each EC number | Database used for homolog search | Number of ESMC layers used for input feature | F1-score |
| --- | --- | --- | --- |
| **1** | **Enzyme-only** | **3** | **0.785** |
| 5 | Enzyme-only | 3 | 0.746 |
| 10 | Enzyme-only | 3 | 0.678 |
| 10 | Enzyme & non-enzyme | 3 | 0.662 |
| 10 | None | 3 | 0.653 |
| 10 | None | 1 | 0.643 |
| 10 | None | 2 | 0.651 |
| 10 | None | 4 | 0.651 |
| 10 | None | 5 | 0.651 |
| 10 | None | 10 | 0.648 |

Second, existing deep learning models usually extract sequence features from the target protein sequence itself. On the other hand, EZpred extracts sequence features from both the target protein and sequence homologs detected by MMseqs2 (17) sequence search. Including sequence homologs results in 2% improvement in the F1-score for EC number prediction. Additionally, including only enzymes in the sequence database for MMseqs2 sequence search results in 4% greater F1-score than including both enzymes and non-enzymes in the database (**Table 1**).

Third, with very few exceptions (34, 51), most existing models that use protein language models for sequence feature extraction only use the output from the last layer. On the other hand, when EZpred uses the ESMC model for feature extraction, outputs from the last three layers of ESMC are all used. The ablation study shows that using these three layers results in 2% and 0.8% greater F1-score than using only the last layer and the last 10 layers (**Table 1**).

Finally, the template-based component of EZpred combines both sequence search and structure search results from MMseqs2 and Foldseek (23), respectively, where the structures of the target and templates proteins are from the AlphaFold protein structure database (AFDB) (52). The combined template-based component achieves an F1-score of 0.785, which is 2% higher than those using sequence and structure hits only (F1-score=0.767), indicating that both sequence and structure templates are helpful for function prediction.

### Case study

As a case study, we investigated the EC number prediction for 7,8-linoleate diol synthase from the fungus *Pyricularia oryzae* (UniProt accession: G4N4J5). This synthase is a bifunctional enzyme with two EC numbers: EC 1.13.11.60 “linoleate 8R-lipoxygenase” and EC 5.4.4.6 “9,12-octadecadienoate 8-hydroperoxide 8S-isomerase”. EC 1.13.11.60 converts linoleic acid (18:2n-6) into 8-hydroperoxy-8(E),12(Z)-octadecadienoic acid (8-HPODE, **Figure 4A**), while EC 5.4.4.6 catalyzes the isomerization of 8-HPODE to 7,8-dihydroxy-9(Z),12(Z)-octadecadienoic acid (7,8-DiHODE, **Figure 4B**). Among the 8 existing predictors, none of them can predict EC 5.4.4.6 for this protein; only two predictors (ProteInfer and DeepEC) can correctly predict EC 1.13.11.60 (**Figure 4C**). On the other hand, both the template-based and deep learning components of EZpred correctly predict EC 1.13.11.60, and the latter also correctly predicts EC 5.4.4.6. Meanwhile, both components make a false positive prediction EC 5.4.4.5 which catalyzes isomerization of 8-HPODE to 5,8-dihydroxyoctadeca-9,12-dienoate (5,8-HiHODE) instead of 7,8-DiHODE (**Figure 4D**). This is due to several Psi-producing oxygenase A enzymes in the training set that performs EC 1.13.11.60 and 5.4.4.5 and have high similarity to the target enzyme (UniProt accessions: G5EB19, Q4WPX2, Q6RET3, **Figure 4E**). These three template enzymes are ranked 3^rd^, 4^th^ and 5^th^ among all templates, leading to the partially incorrect prediction. Nonetheless, these data show that combining template (F1-score=0.667) and deep learning (F1-score=0.500) can lead to more accurate prediction (F1-score=0.800, **Figure 4C**).

**Figure 4.**
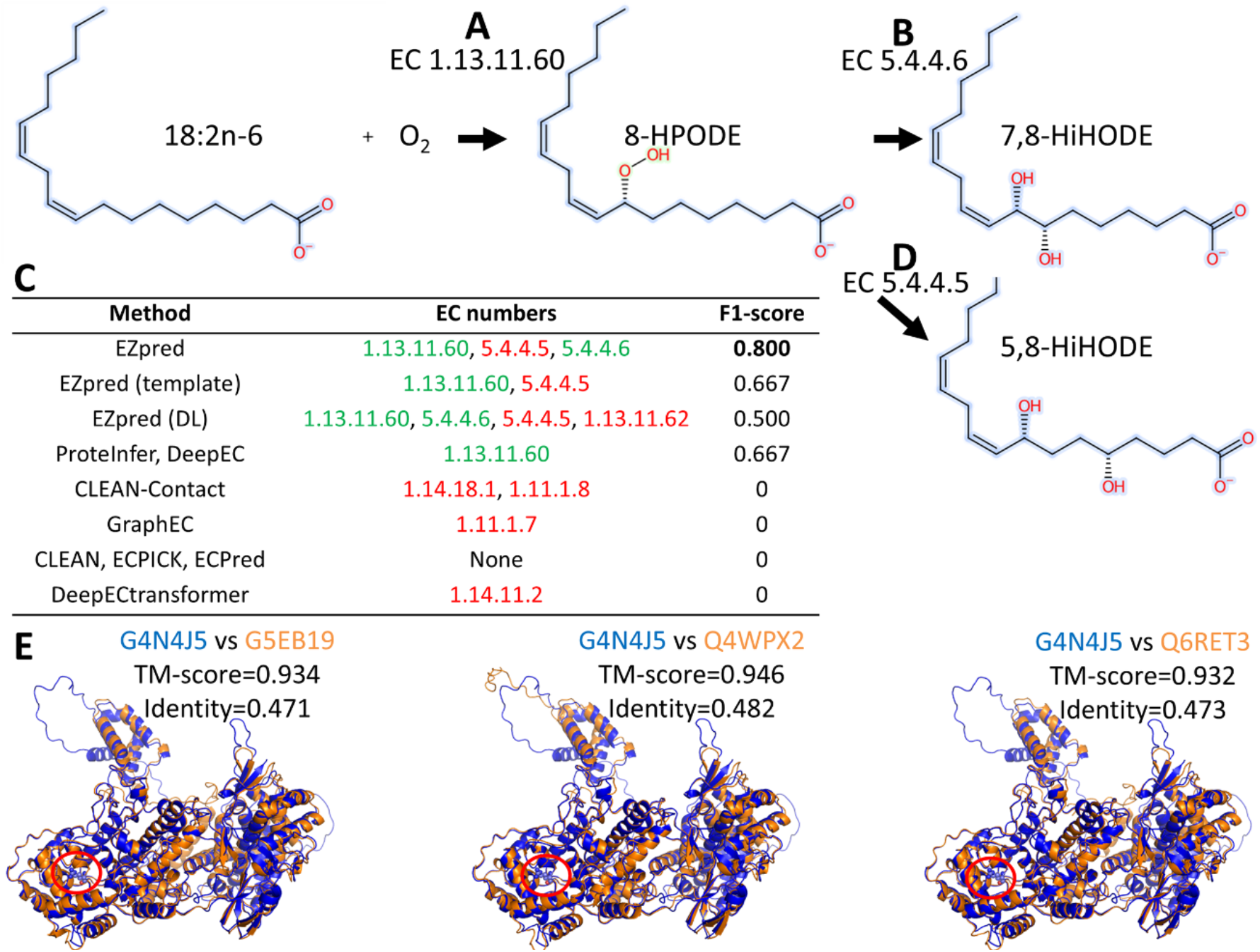
Case study of 7,8-linoleate diol synthase (UniProt accession: G4N4J5). **A**. EC 1.13.11.60 “linoleate 8R-lipoxygenase”. **B**. EC 5.4.4.6 “9,12-octadecadienoate 8-hydroperoxide 8S-isomerase”. **C**. EC prediction by EZpred and existing methods. Green and red indicate correct and incorrect predictions. **D**. EC 5.4.4.5 “9,12-octadecadienoate 8-hydroperoxide 8S-isomerase”. **E**. Structure superimposition between G4N4J5 (blue) and three structure templates (orange). Red circle indicates the catalytic active site.

## Discussion and Conclusion

In this work, we develop EZpred, a composite pipeline that combines deep learning, sequence homolog search and structure template alignment for the prediction of enzyme function. In a large-scale benchmark dataset, EZpred consistently outperforms existing deep learning methods by >27% higher F1-score when predicting the full 4-digit EC number and >17% higher F1-score when predicting whether a protein is an enzyme. Detailed analysis shows that including rare EC numbers, extracting features from sequence homologs, using the last three layers of the ESMC protein language model, and combing sequence and structure templates all contribute positively to the overall performance of EZpred.

Despite the high performance of EZpred, there are still two points that are worthy of further development efforts. First, while its template-based component uses both sequence and structure for template search, EZpred does not use tertiary structure in its deep learning component. Integration of structural features (53, 54) may further improve the deep learning-based component. Second, enzyme function is mainly determined by the active site residues and residues within the substrate/product binding pocket. Yet, the mean pooling procedure in the deep learning-based component treats all residues equally. Therefore, future versions of EZpred will use higher weights for predicted active/binding site residues when pooling features for the deep learning component.

## Materials and Methods

EZpred contains two components: the template-based component and the deep learning-based component (**Figure 1**).

### Template-based component of EZpred

The template-based component of EZpred (red component in **Figure 1**) includes two parts: sequence template search and structure template search.

In the sequence template search step, the input protein sequence is searched by MMseqs2 through all training enzyme protein sequence using the following search parameters: “easy-search --max-seqs 4 --format-output query,qlen,target,tlen,evalue,bits,alnlen,fident,qaln,qstart,taln,tstart -e 1e-2”. For any EC number *q*, its sequence template-based prediction score is calculated as:

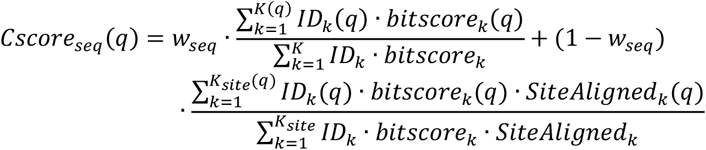

Here, *bitscore*_*k*_ is the bit-score for aligning the target protein to the *k*-th template sequence. *ID*_*k*_ is the global sequence identity for the *k*-th template and equals the number of identical amino acid residues divided by the maximum between the target sequence length and template sequence length. *K* and *K*_*site*_ are the total number of templates and the subset of templates with known active sites or substrate binding sites according to the UniProt database. In UniProt, active sites are annotated by UniProt through the “FT ACT_SITE” tag. Substrate binding sites are also detected from the UniProt data file by the following procedure: ChEBI IDs of the substrates and products of the catalytic activity is extracted from the “CC -!- CATALYTIC ACTIVITY” records; second, ligand binding residues are detected from the “FT BINDING” and “FT REGION” records; finally, the subset of binding residues associated with ChEBI of the substrate/product molecules via the ‘FT /ligand_id=“ChEBI:CHEBI:”)’ records are confirmed to be substrate binding residues. We then defined the catalytic site pocket by including all active site residues, substrate binding site residues, and all other residues whose sequence separations to the above residues are ≤2. *SiteAligned*_*k*_ is the portion of catalytic site pocket residues in the *k*-th template that are aligned to the target protein. Meanwhile, *bitscore*_*k*_*(q), ID*_*k*_(q), *K*(q), *K*_*site*_ (q), and *SiteAligned*_*k*_*(q)*are for the subset of templates with EC number *q*. The weighting between global and local similarity, *w*_*seq*_, is defined as:

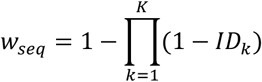

In the structure template search step, the input protein structure is searched by Foldseek through all training enzyme protein structures from AFDB using the following search parameters: “easy-search --format-output query,qlen,target,tlen,evalue,bits,alnlen,nident,qaln,qstart,taln,tstart, qtmscore,ttmscore --tmalign-fast 1 -e 0.1 -s 7.5”. For EC number *q*, its structure template-based prediction score is calculated as:

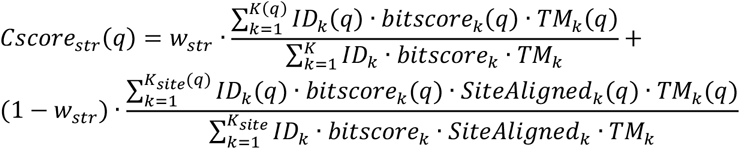

Here, *TM*_*k*_ is the TM-score between the target protein and the k-th template protein. *w*_*str*,_ uses the same formula as *w*_*seq*_, except that the sequence identities of the Foldseek-detected structure templates rather than those of the MMseqs2-detected sequence homologs are used for its caclulation. If the target and template proteins have different lengths, there will be two TM-score values corresponding to the two sequence lengths, in which case the minimum between the two TM-score values is used. The sequence and structure template-based scores are then combined linearly:

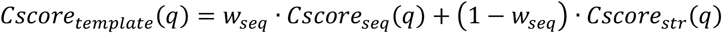

### Deep learning-based component of EZpred

The deep learning-based component of EZpred (blue component in **Figure 1**) is inspired by our previous pipelines InterLabelGO (51) and ATGO (34). This component starts with feeding the target protein sequence into the ESMC protein language model to generate a sequence embedding matrix from its last three hidden layers. Each layer produces a 1152 × *L* matrix, where *L* is the length of the target protein. The ESMC model can only process up to 2048 tokens. Thus, if *L* > 2048, only the first 2048 residues of the protein sequence are considered. Mean pooling is applied along the sequence to generate an embedding vector with 1152 elements. These three vectors from mean pooling of the last three layers of ESMC are concatenated into a single embedding vector with 1152×3 = 3456 elements.

Meanwhile, the target protein is also searched through the sequence database consisting of training enzymes using MMseqs2 to identify sequence homologs (See above Methods section for template-based component of EZpred). Only the top four MMseqs2 hits are considered. Each of these four sequence homologs are passed through ESMC, and the embeddings from the last three layers of ESMC are mean pooled into an embedding vector with 3456 elements as above. These four embedding vectors, denoted as 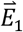 to 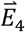, are combined into a single embedding vector 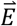 with 3456 elements by the following formula weighted averaging:

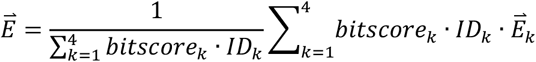

The combined embedding vector 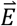 and the embedding vector from the target protein itself are concatenated into a single embedding vector with 3456 × 2 = 6912 elements. It is further processed by a feed forward neural network with the Gaussian Error Linear Unit (GELU) activation function to transform the ESMC-derived features into EC number probability. Based on predicted labels, we train two types of neural network models. The first type of model predicts 928 EC numbers, including 7 EC numbers with 1 digit, 64 EC numbers with the first 2 digits, 188 EC numbers with the first 3 digits and 665 EC numbers with all 4 digits. The second type of model predicts all the above EC numbers as well as an additional label (0.-.-.-) for non-enzyme proteins. Since we use 5-fold cross validation, we generate 5 models for each model type. The prediction scores for EC numbers and for non-enzymes are derived from averaging the prediction result from the 5 models of the first type and second type, respectively. We use a dropout rate of 30% and a batch site of 512. Early stopping is applied if a model is trained with at least 20 epochs and the validation loss has not been improved for the last 10 epochs.

Similar to InterLabelGO, the neural network of EZpred is trained using a composite loss function consisting of the protein-centric F1-score, the EC number-centric F1-score, and the Zero-bounded Log-sum-exp and Pairwise Rank-based (ZLPR) loss (55). Since the differences in the probability of different EC numbers are large, we use information content (IC, also known as information accretion) (56) to weigh different EC numbers, where the IC for EC number *q* is defined as:

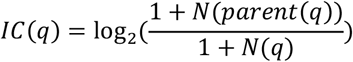

Here, *N*(*q*) is the total number of training enzymes with EC number *q*; *parent*(*q*) is the parent of EC number *q*, i.e., the EC number with one less digit. For example, for EC number 2.7.4.3 “Adenylate kinase”, its parent EC number is 2.7.4.- “Phosphotransferases with a phosphate group as acceptor”. If *q* only has one digit, *N*(*parent*(*q*)) equals the number of all training enzymes. Using the IC, we define the precision and recall for protein *t*:

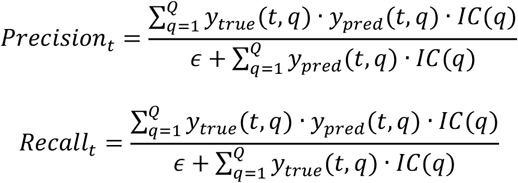

Here, *ϵ*=1E-16 is a small value to avoid division by zero; *y*_*true*_(*t, q*) indicates whether protein *t* has EC number *q*; and *y*_*pred*_ (*t, q*) is the predicted probability of EC number *q* for protein *t*. Q=928 is total number of labels, i.e., EC numbers. Similarly, the precision and recall can also be defined for EC number *q*:

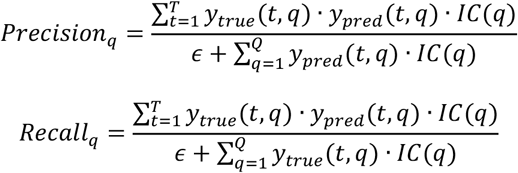

Here, *T* is the total number of training proteins. The protein-centric F1-score loss can be defined as:

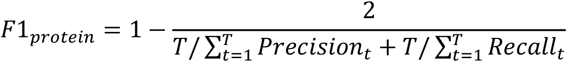

Similarly, the EC-centric F1-score loss can be defined as:

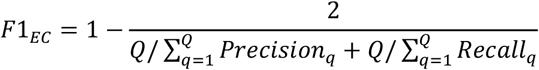

The third term for the loss function is the ZLPR loss (55), which was found in our previous study (51) to be a more effective replacement compared to the traditional binary cross entropy loss. The ZLPR loss is defined as:

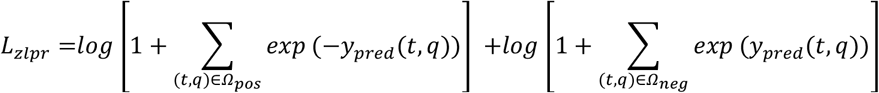

Here, *Ω*_*pos*_ and *Ω*_*neg*_ are the sets of positive and negative protein-EC number associations. The final composite loss function used to train EZpred is:

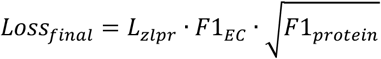

To ensure the true-path rule, i.e., that the predicted probability of EC number *q, Cscore*(*q*), is no greater than the predicted probability of parent EC number *Cscore*(*parent*(*q*)), and no less than the maximum predicted probability of any of its child EC number *Cscore*(*child*(*q*)), an iterative post-processing procedure is applied. In each iteration, the following steps are applied. If *Cscore*(*parent*(*q*))≥*Cscore*(*q*)≤*Cscore*(*child*(*q*)), *Cscore*(*q*) is increased to max(*Cscore*(*parent*(*q*)), *Cscore*(*child*(*q*))). If *Cscore*(*parent*(*q*))≤*Cscore*(*q*)≥*Cscore*(*child*(*q*)), *Cscore*(*q*) is decreased to min(*Cscore*(*parent*(*q*)), *Cscore*(*child*(*q*))). If *q* has all 4 digits and no child EC number and *Cscore*(*parent*(*q*)) < *Cscore*(*q*), *Cscore*(*q*) is decreased to the value of *Cscore*(*parent*(*q*)). If *q* has only 1 digit and no parent EC number, and *Cscore*(*q*)<*Cscore*(*child*(*q*)), *Cscore*(*q*) is decreased to the value of *Cscore*(*child*(*q*)). If *Cscore*(*parent*(*q*))<*Cscore*(*q*)<*Cscore*(*child*(*q*)), *Cscore*(*q*) is not updated in the current iteration. The above iteration is performed up to 4 times, until the predicted probability of any EC number is no greater than that of its parent.

The template-based score *Cscore*_*template*_*(q)*is then combined with the deep learning-based prediction *Cscore*_*DL*_*(q)*to derive the final EZpred prediction score for EC number *q*:

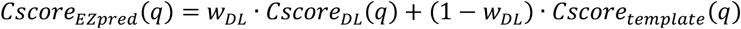

The weight to balance the two components equals to *w*_*DL*_ = 0.2 +0.1·(1−*w*_*seq*_). Here, the values 0.2 and 0.1 are hyperparameters optimized by a grid search through cross-validation.

## Acknowledgments

The authors thank Dr Xiaoqiong Wei for insightful discussions. This work used the Advanced Cyberinfrastructure Coordination Ecosystem: Services & Support (ACCESS) program, which is supported by National Science Foundation (2138259, 2138286, 2138307, 2137603, and 2138296). This work is supported in part by the National Institute of Allergy and Infectious Diseases (AI134678 to L.F).

